# Sawfish, Read in Tooth and Saw: rostral teeth as endogenous chemical records of movement and life-history in a critically endangered species

**DOI:** 10.1101/753293

**Authors:** Jens C. Hegg, Breanna Graves, Chris M. Fisher

## Abstract

1. The ecology of endangered and rare species can be difficult to study due to their low abundances and legal limits on scientist’s ability to catch, sample, and track them. This is particularly true of sawfish (family Pristidae) whose numbers have declined precipitously, placing all five species on the IUCN Red List of Threatened Species worldwide. Best known for their distinctive, toothed rostrum, the ecology, movement, and life-history of sawfish is poorly understood.
2. Sawfish rostral teeth are modified placoid scales, which grow continuously throughout the life of the fish. This continuous growth, combined with their stable calcified makeup, makes sawfish rostral teeth a potential source of temporal records of chemical and isotopic changes through the life of the fish.
3. Rostral teeth are often preserved in museums and as curios, potentially providing a source of life-history data to inform conservation actions without the need for field study, or as an important compliment to it. This is the first study to recover temporally explicit chemical data from sawfish rostral teeth.
4. Using archived samples of largetooth sawfish (*Pristis pristis*) we show that multiple chemical tracers can be recovered from sawfish rostral teeth, and that these tracers can be used to understand movement across salinity gradients. We further show that sawfish rostral teeth contain repeated structures and indistinct banding which could potentially be used for aging or growth analysis of fish.

## 1. Introduction

Understanding the ecology of endangered and cryptic species can be challenging. Low abundance, and the ethical limitations on human interaction for highly endangered species, limit our ability to apply traditional techniques to understand the ecology of these species at appropriate temporal and spatial scales (Cuthill, 1991; Mackenzie et al., 2005; Chadès et al., 2008; Parris et al., 2010). However, effective conservation of these species often requires detailed understanding of their life-history, migration patterns, and ontogeny. This leaves ecologists to find creative ways to uncover this important information.

The largetooth sawfish (*Pristis pristis*), a large ray with an iconic toothed rostrum, is a critically endangered species with extremely low abundances whose life-history is poorly understood (Burgess, Fernandez-Carvalho & Imhoff, 2009; Kyne, Carlson & Smith, 2013; Fernandez-Carvalho et al., 2014). Along with habitat loss and black market trade in fins and rostra, a major threat to sawfish is entanglement in nets as bycatch for other species (Seitz & Poulakis, 2006; Dulvy et al., 2016). All species of sawfish move across a range of salinities from salty to brackish, and largetooth sawfish, are found in salty and fresh water (Thorson, 1982; Poulakis et al., 2011; Scharer et al., 2012; Faria et al., 2013).

Behavior of Australian largetooth sawfish populations in relation to salinity are relatively better understood (Peverell, 2005; Thorburn et al., 2007; Whitty et al., 2009; Gleiss et al., 2017). However, regional variation in freshwater use in largetooth sawfish is as evidenced by historical populations in Lake Nicaragua and reports as far upstream as Manaus in the Amazon river indicating extensive freshwater use in some populations (Thorson, 1982; Burgess, Fernandez-Carvalho & Imhoff, 2009). Despite this, the ecology and distribution of largetooth sawfish in Central and South America is not well studied and many published accounts are limited to opportunistic sightings (Nunes et al., 2016; Feitosa, Martins & Nunes, 2017; Schmid & Giarrizzo, 2017). However, understanding the distribution and ecology of sawfish, especially in their remaining stronghold in the Amazon region, is a necessary precursor to conservation actions beyond the legal limits on catch and trade in sawfish parts (Kyne, Carlson & Smith, 2013; Dulvy et al., 2016). Uncovering patterns of habitat use and migration, and their relationship to net fishing which is a large source of mortality, could help target conservation and management actions to life-stages, locations, or seasons where the risk to sawfish is greatest. Further, understanding the locations of nursery or rearing areas in fresh and brackish water could promote conservation of these locations. However, low population densities and large expanses of potential habitat make collecting movement data using capture or tracking technologies particularly difficult.

Sawfish conservation across a large portion of their range is limited by a lack of knowledge related to their presence or absence, and the location of critical habitats (Dulvy et al., 2016). This lack of specific spatial knowledge, and the reality of extremely low catch rates, is reflected in recent innovative use of indirect abundance-occupancy models to assess the likelihood of sawfish abundance and occupancy worldwide (Yan et al., 2021). The presence of sawfish in much of Latin America, Africa, and Asia remains listed as “uncertain” or “possibly extinct” for multiple species (Dulvy et al., 2016). The range of the most widely distributed species, the largetooth sawfish, has primarily been defined by historical accounts across much of their range, with the vast majority of our detailed knowledge coming from Lake Nicaragua and Northern Australia, and even stongholds in the Amazon region have not been systematically surveyed (Kyne, Carlson & Smith, 2013; Kyne et al., 2021). Further, the opportunity to collect new information in many areas is considered low, due to this decrease in population abundance (Kyne et al., 2021).

Where detailed presence or absence data is missing, conservation plans are often built based upon community knowledge of sawfish catches, with opportunistic collection of size and catch data where samples are available (Leeney & Poncelet, 2015; Jabado et al., 2017; Mendoza et al., 2017; Haque, Leeney & Biswas, 2020; López-Angarita et al., 2021). While historical data may not reflect the current state of the population, it increasingly plays a role in planning and management within marine science, as conservation often requires an understanding of both historical range and current information to define conservation goals (Máñez et al., 2014). However, community-based approaches and opportunistic catch data is often of low-resolution. Further, designation of important habitat is often done without the opportunity for thorough assessment of local behavior and ecology, as evidenced by habitat designation of the smalltooth sawfish in the United States which is only now being supplemented by direct studies of movement and location using telemetry (Wiley & Simpfendorfer, 2010; Norton et al., 2012; Graham et al., 2020), and presence/absence studies using environmental DNA (eDNA) techniques (Simpfendorfer et al., 2016; Lehman et al., 2020).

In areas where funding is available for systematic and large-scale population research, telemetry and eDNA data is extremely useful. However, both methods require significant funding and investment, funding which is often unavailable in much of the sawfishes range. Further, these methods can only reveal the movements and location of a severely depleted species which has lost significant range, and presumably similarly significant levels of life-history diversity.

The importance of habitat and life-history diversity on the resilience of species, and the importance of identifying and prioritizing life-history diversity within conservation plans is well understood (Beechie et al., 2006; Schindler et al., 2010; Schroeder et al., 2016). New tools which can increase the data richness of historical samples are needed, and attempts are currently underway to use archived rostral samples to understand historical genetic diversity and population structure of sawfish worldwide (Phillips, Nicole., Wueringer, 2015; Fearing et al., 2018). However, no tools currently exist which can elucidate life history in these archived rostral samples, or tie historical genetic information gleaned from them to the movement and behavioral aspects of sawfish ecology. Further, few opportunities exist to determine adult life-history of sawfish, one of the least understood areas of sawfish ecology (Kyne et al., 2021), yet many archived rostrum represent samples of large adult fish.

One tool that has been revolutionary in understanding movement and ecology of teleosts (and increasingly chondrichthyans) is the use of chemistry stored in fish hard parts to determine provenance, migration, and other aspects of life history (Kennedy et al., 1997; Hobson, Barnett-Johnson & Cerling, 2010; Tillett et al., 2011; Smith, Miller & Heppell, 2013; Walther, 2019; Wang, Walther & Gillanders, 2019). These isotopic and trace elemental techniques are particularly powerful when applied to calcareous structures which grow throughout the life of the fish, as other Group 2 elements readily substitute for calcium in the matrix, and other elements are incorporated in the crystalline structure (Elsdon et al., 2008).

Strontium isotope composition, ^87^Sr/^86^Sr, has been shown to be recorded without fractionation in fish otoliths (Kennedy et al., 1997, 2000). This isotope ratio can vary greatly in nature and has been used to trace fish movement with kilometer precision in fresh water, and the presence of a global marine signature worldwide can be used as a marker of entry into salt water (Hamann & Kennedy, 2012; Courter et al., 2013; Hegg et al., 2013).

The ratio of elemental Sr and Ba to calcium (Sr/Ca and Ba/Ca) in hard parts have been shown to be useful tracers of salinity changes due to generally higher bulk strontium in the ocean than fresh water, and the flocculation of barium in the presence of salinity (Kalish, 1990; Zimmerman, 2005; Shippentower, Schreck & Heppell, 2011; Hamer et al., 2015). This has been used to understand entry of anadromous fish into salt water, as well as habitat use in estuarian and euryhaline species, and to distinguish provenance of fully marine coastal species as well (Kraus & Secor, 2004; Warner et al., 2005; Mohan et al., 2012; Schaffler, Miller & Jones, 2014; Rohtla & Vetemaa, 2016; Seeley & Walther, 2017). As well, trace element ratios of Mg (Mg/Ca) have been strongly related to growth and metabolism with potentially higher influx into otolith endolymph through ion channels during higher metabolic activity (Limburg et al., 2018). Manganese is rarely found in the water column in an ionic form suitable for incorporation, and is released only in or near reducing, anoxic substrates (Altenritter, Cohuo & Walther, 2018; Limburg & Casini, 2018). Therefore, Mg/Ca has been linked to exposure to reducing environments such as mangroves, and exposure to hypoxic stress in several species has been well described (Mohan et al., 2014; Paillon et al., 2014; Turner, Limburg & Palkovacs, 2015; Smith et al., 2016; Altenritter, Cohuo & Walther, 2018; Limburg & Casini, 2018). Further, numerous studies in anadromous, catadromous, coastal and estuarine species utilize a suite of these elements and isotopes to determine provenance. These include tracing provenance and of reef fish to natal seagrass beds, natal origin of American shad across the Atlantic seaboard, nursery areas of menhaden in Chesapeake Bay, entry of juvenile salmon into the ocean, and facultative oligohaline habitat use in Atlantic tarpon (Chittaro et al., 2006; Walther & Thorrold, 2008; Hegg, Kennedy & Fremier, 2013; Schaffler, Miller & Jones, 2014; Woodcock & Walther, 2014).

Until recently the use of chemical signatures in sharks and rays has lagged behind their use in teleost species and has been confined mainly to elemental ratios due to the utility of elemental ratios in fully marine species. However, recent studies have shown that elasmobranch vertebra function as temporal records of elemental chemistry and location (Tillett et al., 2011; Smith, Miller & Heppell, 2013; Raoult et al., 2016). Numerous studies have now been published using suites of elemental markers to determine location and stock structure in elasmobranchs (Izzo et al., 2016; McMillan et al., 2017; Mohan et al., 2018; Coiraton, Amezcua & Ketchum, 2020). Further work utilized strontium and barium elemental ratios to recover movements across salinity gradients in several species of shark, including repeated spawning movements into fresh water for bull shark, and likely proximity to mangrove areas based on manganese concentrations (Tillett et al., 2011; Lewis et al., 2016; Feitosa, Dressler & Lessa, 2020). Recent work has shown a similar relationship of salinity to Sr/Ca in smalltooth sawfish vertebra as that shown in other elasmobranchs and teleosts (Scharer et al., 2012).

The incorporation of these elemental signatures into elasmobranch hard parts appears to function in similar ways to teleosts, though fewer controlled studies have been done. Two controlled tank and mesocosm experiments have shown Sr/Ca, Ba/Ca, Mg/Ca, and Mn/Ca to be similarly useful as natural tags in shark vertebra along with other elemental ratios (Tillett et al., 2011; Smith, Miller & Heppell, 2013). These studies indicate that strontium and barium are taken up in proportion to the element’s concentration in the environment, while elements such as Mg and Mn are more highly regulated at the cellular level and therefore reflect growth and physiological conditions to a greater degree. The effect of temperature on Sr and Ba incorporation in vertebra is conflicting, as it is in the teleost literature, however the magnitude of the temperature effect is small (Tillett et al., 2011). Turnover rate of these signatures within the elasmobranch body is unknown and is likely mediated by the blood supply and speed of growth in the tissue being examined, as opposed to teleost otoliths which lack blood supply but show changes at a scale of weeks (Campana, 1999; Hegg, Kennedy & Chittaro, 2018).

We propose that sawfish rostral teeth may provide a useful record of age, growth and environmental chemistry which could hold information on life-history and movement in individual fish. Rostral teeth are not true teeth but instead highly modified dermal denticles which grow through the life of the fish (Welten et al., 2015). Made of bio-apatite (Miller, 1974; Shellis & Berkovitz, 1980), rostral teeth grow continuously from their base and are not replaced if they are lost (Slaughter & Springer, 1968; Wueringer, Squire & Collin, 2009). Rostral teeth presumably incorporate similar trace-elements and isotopes in their chemical matrix similar to other calcium carbonate and calcium phosphate hard parts such as shells, true teeth, and otoliths (Campana & Thorrold, 2001; Schöne et al., 2004; Begg et al., 2005; Thornton, 2011). While the morphology of sawfish teeth varies across species, their basic formation and growth is similar. These characteristics indicate that sawfish rostral teeth may store useful ecological data as an endogenous record.

Importantly, sawfish rostra are often dried and kept as curios and many are kept in museums, academic and private collections worldwide (Seitz & Poulakis, 2006; Melo Palmeira et al., 2013). These collections provide a reservoir of samples by which to uncover details of sawfish life-history without harming living individuals (Phillips et al., 2009; Whitty et al., 2014). While work from archived collections are necessarily reconstructions of the past, archived samples also potentially reflect records of behavior and life history before and during the ongoing declines in abundance of these species. Because of this, they may hold important information that is no longer evident in extant populations. Further, there is additional potential to utilize natural mortalities and samples seized by law enforcement to infer limited information about current populations.

Only one unpublished study has investigated the use of sawfish rostral teeth as an endogenous record of growth and provenance (Field, Meekan & Bradshaw, 2009). This work indicated that the trace element chemistry of whole sawfish rostral teeth could be used to distinguish sawfish populations around the coast of Australia. The authors also demonstrated that growth banding could be observed on the outside of whole rostral teeth which could be used to estimate age. Despite this, no work has been published investigating whether rostral teeth record higher-resolution chemical or structural data throughout the life of the fish. However, Sawfish vertebra, much like those of sharks and other rays, have been shown to contain yearly growth bands as well as a chemical record of salinity (Scharer et al., 2012).

In this study we present data from the analysis of rostral teeth taken from two largetooth sawfish rostra housed in a university collection in the United States. Our first objective was to demonstrate methods of sectioning, treating and imaging sawfish rostral teeth to quantify age and growth information they contain. Our second objective was to document the potential of sawfish rostral teeth as a chemical record of location and movement using laser ablation-inductively coupled plasma mass spectrometry (LA-ICPMS) of ^87^Sr/^86^Sr isotope ratio, as well as trace element ratios of Sr/Ca, Ba/Ca, Mn/Ca, Mg/Ca. Finally, we analyzed these chemical and growth data to investigate their utility in understanding lifetime sawfish movement across salinity and the existence of other environmental or ontogenetic information recorded during the life of the fish.

## 2. Methods

### Sample Information

All data in this study were taken from two sawfish rostra included in the ichthyology teaching collection at University of Idaho in Moscow, ID USA (Figure 1). The first was the rostrum of a largetooth sawfish embryo. Information attached to the rostrum in the collection read, “Mother caught off Belize (British Honduras) coast, 1969. 8ft long with 5 embryos ≈ 1.5 ft long. Blade covered with protective sheath when embryo was removed from mother.” Thus, the total length (TL) of the embryo was approximately 45 cm. This rostrum has a total rostral length (TRL, length from tip of the rostrum to the attachment to the head where the rostrum begins to flare *sensu* (Whitty et al., 2014)) of 14.2 cm assuming the rostrum was removed at this attachment point, and a standard rostral length (SRL, length from tip of rostrum to the most distal tooth) of 13.6 cm (Figure 1A)

**Figure 1.**
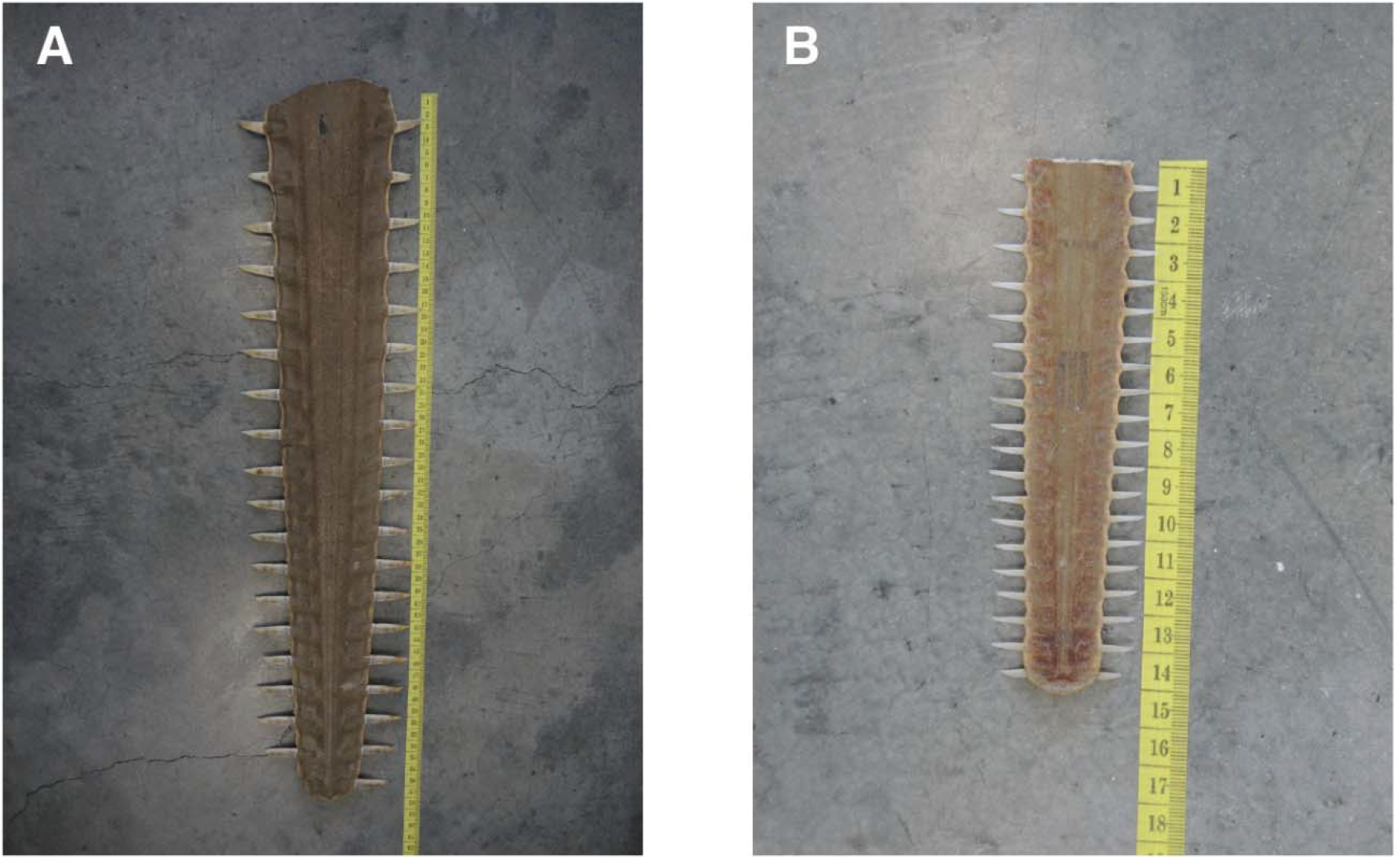
Rostra from two largetooth sawfish were used in this study. A sub-adult rostrum (A) of unknown origin had a TRL of 57cm and a SRL of 55cm. The embryo rostrum collected in Belize in 1969 had a TRL of 14.2cm and a SRL of 13.6cm. The most distal tooth on the left-hand side was taken for analysis from each rostrum.

The second sample is a larger rostrum from a largetooth sawfish with a TRL of 57 cm and a standard rostral length of 55 cm (Figure 1B). No information was available for the provenance of this rostrum. Using the mean SRL to total length (SRL/TL) relationship from Whitty et al. (2014) results in a TL of 238.7 cm for the fish from which the rostrum was taken. This length would indicate the fish was a sub-adult as 240 cm is the smallest published length of maturity for males, with 280 cm for males and 300 cm for females being more widely accepted (Wueringer, Squire & Collin, 2009; Dulvy et al., 2016).

### Structural Analysis

The most distal tooth on the left-hand side of each rostrum was removed using a utility knife and a Dremel tool. The embryo tooth showed no signs of tooth wear. The sub-adult tooth was highly worn on the anterior edge of the tooth extending beyond the rostrum, and significantly pitted and eroded on the posterior edge. Rostral teeth were cleaned of tissue by soaking in water followed by scraping with a scalpel blade. Prior to sectioning, the surface of each tooth was stained using silver nitrate to highlight growth ridges as described by Field et al. (2009). The structure of sawfish teeth is extremely dense, and as noted by Bradford (1957) is difficult even to dissolve using nitric acid. Therefore, this surface treatement is unlikely to affect further chemical analysis within the sectioned tooth.

Rostral teeth were then embedded in epoxy resin and sectioned using an Isomet low speed saw and a diamond wafering blade (Beuhler, beuhler.com) in a manner described for large otoliths and fin rays (Secor, Dean & Laban, 1992; Koch & Quist, 2007). The sub-adult tooth was curved both distally and along the dorsal-ventral axis. To allow sectioning through the center of the entire tooth, given the complex curvature, it was cut in half near the point at which it protruded from the rostrum. The embryo tooth and the two pieces of the sub-adult tooth were embedded in epoxy and an approximately 1mm section in the dorsal-ventral plane was taken that included the center core of each tooth (Figure 2). A further two sections were taken in the same plane from the sub-adult tooth, and a second section was taken from the embryo tooth to facilitate staining while maintaining one unstained section for chemical analysis. Each section was polished on 3600 grit and 6000 grit Micro Mesh wet-dry sandpaper (Micro Mesh, micromesh.com).

**Figure 2.**
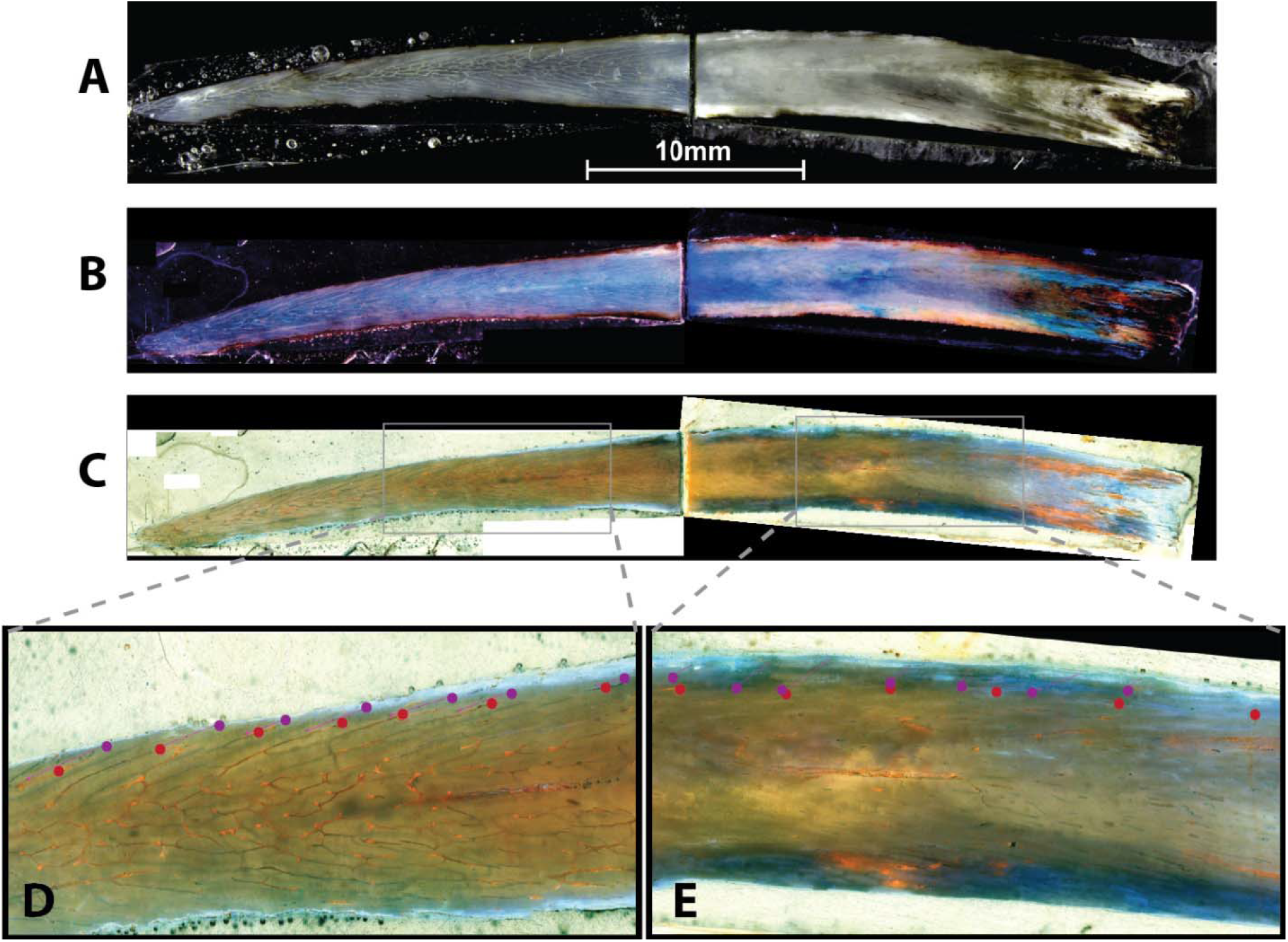
Sub-adult largetooth sawfish rostral tooth sectioned on the dorsal-ventral plane (A) shows internal tubule structures and faint banding. Staining with Mutvei’s solution (acetic acid, glutaraldehyde, and alcian blue) highlighted the tubules which broke the surface in darker blue (B). Inverting the color spectrum of the stained section (C) further improved visualization of the surface tubules. Tubules overlapped in curved bands (D), with “legs” descending distally from the core of the tooth to the edge. These tubules were less distinct near the base of the tooth, however faint banding was also visible. Putative growth was quantified by two readers (reader 1 – red dots, reader 2 – purple dots). Measurements agreed despite being offset where descending tubule “legs” were prominent (D). Agreement was lower in sections with less prominent tubules (E).

The sub-adult tooth sections were stained using both silver nitrate and Mutvei’s solution (Dunca, Fiebig & Pfeiffer, 2005) to enhance any growth banding present in the rostral teeth. Sections were submerged in silver nitrate for approximately 5 minutes prior to light exposure. The second section was submerged in Mutvei’s solution (dilute acetic acid, gluteraldahyde, and alcian blue) for 6 hours. The remaining embedded sample was burned using a handheld butane micro-torch (BernzOmatic, bernzomatic.com) until the sample surface was golden brown. The sample was then polished using the same sequence of wet-dry sandpaper to enhance the rings. The embryo tooth was not stained for growth analysis as the small size of the tooth caused the second thin section to be non-representative of the core of the tooth and very little internal structure was present to indicate growth analysis would be fruitful. Therefore, we opted to preserve the best thin section for chemical analysis.

Each section was imaged in sections at 10X magnification using a Leica DFC450 digital camera (leicacamerausa.com) attached to a Zeiss AxioImager A.1 microscope (zeiss.com). These images were subsequently stitched into a larger mosaic using the Photomerge tool in Adobe Photoshop (adobe.com) with the “collage” layout option and “blend images together” option checked. These color images were then transformed into grayscale and the color palette was inverted to determine an imaging method which would best highlight any banding or sequential growth structures that were present. Each image was subsequently optimized for contrast using a levels adjustment.

The number of bands and/or sequential structures inside the sub-adult tooth were counted by two readers using ImageJ analysis software (https://imagej.nih.gov). Points and distances between them were measured along a line adjacent to the proximal edge of the tooth, on the outside of the dentin layer and inside the enamel layer where it was still present. Distances were standardized to the length of the laser scan for comparison with the chemical data.

### Trace-element and Isotopic Analysis

The unstained sections of the tip and base of the sub-adult rostral tooth, as well as the embryo, were analyzed for strontium composition (^87^Sr/^86^Sr) as well as concentration of a suite of trace elements (^86^Sr, ^138^Ba, ^55^Mn, ^25^Mg). The structure of sawfish teeth is dentin, primarily composed of calcium phosphate (Miller, 1974; Shellis & Berkovitz, 1980). Thus, calcium was used as the internal standard to correct for variation in laser ablation efficiency across the sample and all trace element data is expressed as a ratio to calcium (Sr/Ca, Ba/Ca, Mn/Ca, Mg/Ca).

Analysis was conducted at the Radiogenic Isotope and Geochronology Laboratory (RIGL) at Washington State University. Strontium ratio and trace element data were collected simultaneously using a laser ablation split stream (LASS) methodology (Hegg & Fisher, 2020). Laser ablation was conducted using a Teledyne Analyte Excite 193nm ArF laser using a 35μm spot diameter, 10μm/sec rastering speed, operating at 20Hz, and with laser fluence of ∼4J/cm^2^. Ablated material was then split between a NeptunePlus multicollector inductively coupled mass spectrometer (MC-ICPMS) for ^87^ Sr/ ^86^Sr measurement and Element2 ICPMS for trace element measurement. The laser was used to sample along the center of the sampled tooth, perpendicular to the growth axis, from the tip to the base of the tooth.

Trace element concentrations were normalized to the NIST-610 glass standard, while ^87^Sr/ ^86^Sr was corrected for K and Rb interferences and normalized to a modern marine shell and a ^87^Sr/ ^86^Sr of 0.70918 for modern ocean water, as described by (Hegg & Fisher, 2020). Trace element analysis was calibrated using the NIST 610 standard as described in (Hegg & Fisher, 2020).

All data reduction was completed using a custom built data reduction reduction scheme (DRS) operating within the Iolite software package (Fisher et al., 2017; Hegg & Fisher, 2020). All plotting and analyses were completed in R statistical software (https://www.r-project.org).

## 3. Results

### Structural Analysis

Whole rostral teeth, both unstained and after silver nitrate staining, did not exhibit the external growth banding described by Field et al. (2009). The interior of both rostral teeth contained an inner, clear layer and an outer opaque layer in areas which had not been subjected to wear (Figure 2A). The rostral teeth contain a mesh-like network of interconnected tubules as described by Miller (Miller, 1974). These tubules followed a general pattern of intertwining tubes near the central core, with downward “legs” curving distally toward the edge of the tooth (Figure 2A&D). These descending legs appear to be spaced somewhat regularly, suggesting that they could mark growth or age.

Burning of the sample showed some evidence of layering within the tooth but it was too indistinct to be easily quantified. Silver nitrate staining of the tooth sections did not reveal additional layering or banding structure. Mutvei’s solution was moderately successful in bringing out faint alternating banding in the middle portion of the sub-adult tooth (Figure 2B, D&E). The alcian blue stain in Mutvei’s solution infiltrated the tubules which broke the surface of each tooth section, making clear which tubules were on the surface and which were below.

Both readers agreed that the inverted image of the section stained with Mutvei’s solution provided the best image for measurement (Figure 2C). This image rendered the surface tubules in a distinct orange which was easier to discern than the blue of the non-inverted image. Quantifying of the apparent banding created by the tubule legs and indistinct light-dark bands was done by both readers. Both readers picked 26 growth bands. The first 15 of these bands, starting from the youngest growth at the tooth tip, matched between the readers (Figure 2D). In the basal section of the tooth, where the tubules were most indistinct, 6 of the remaining 11 points matched between the readers (Figure 2E). A consensus measurement was then created which contained 29 bands. When plotted the distance between measurements displayed a linear relationship.

### Trace-element and Isotopic Analysis

Laser ablation and ICP-MS analysis was successful, with 35μm spot size allowing useful resolution across the sample (Figure 3). The marine shell standard measured 0.70915 (±0.00038 2SE, global marine signature 0.70918). Trace element measurements exceeded the limits of detection (LOD) by at least three orders of magnitude in all cases (Average LOD across standards and samples: Sr=0.24 ppm, Ba=0.07 ppm, Mn=0.36, ppm, Mg=0.54 ppm). All analyses exhibited repeated, low level, spikes in signal. While tubules at the base of the rostral tooth created voids which unavoidably resulted in large changes to the signal, the smaller, repeated spikes appeared not to correlate with voids in the surface of the tooth or with changes in the signal and were therefore assumed to be environmental.

**Figure 3.**
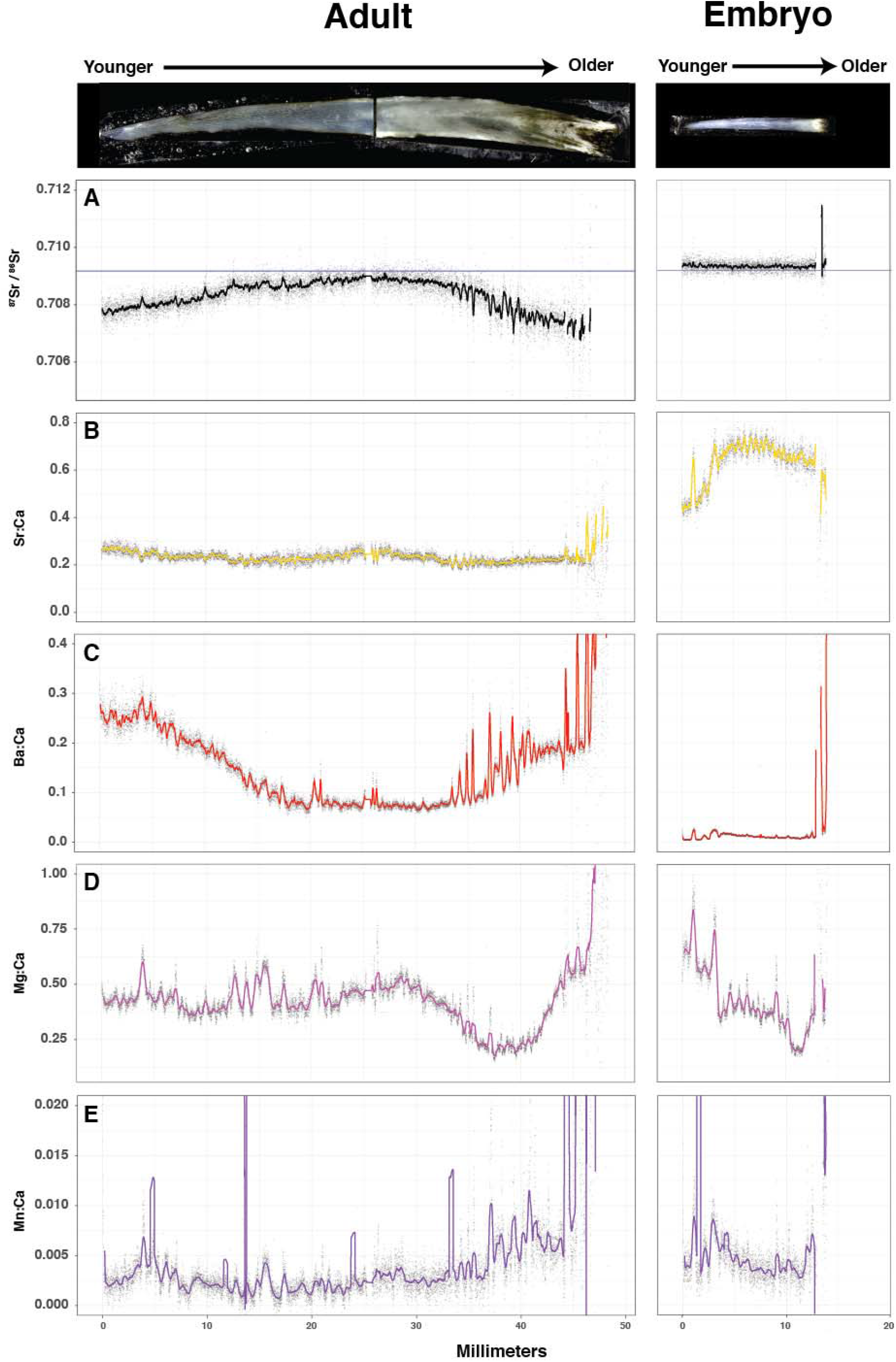
Results of split stream LA-ICPMS analysis of sectioned rostral teeth from to largetooth sawfish specimens, sub-adult (left-hand column) and an embryo (right-hand column) showed significant variation across the tooth. Relative age of the fish is indicated above the plots, with the youngest age represented by zero microns, and older ages with increasing distance along the tooth. Strontium ratio (^87^Sr/^86^Sr, A) indicated movement within fresh or brackish water, with movement toward the global marine signature (blue horizontal line) in mid-life indicating movement into higher salinity water. Meanwhile the embryo ^87^Sr/^86^Sr appears to match the global marine signature throughout life. Strontium to calcium ratio (Sr/Ca, B) did not rise significantly during mid-life in the sub-adult, confirming residence in brackish rather than fully saline water. The embryo Sr/Ca, in contrast, is elevated, confirming a likely ocean residence during provisioning or development. Barium to calcium ratios (Ba/Ca, C) increase in fresh water and decrease quickly in the presence of salinity, confirming sub-adult and embryo movements inferred from the prior tracers. Magnesium to calcium ratios (Mg/Ca, D) are linked to growth rate in other species. Peaks in Mg/Ca could indicate growth rate changes as well as movement into new locations. Manganese to calcium ratios (Mn/Ca, E) are linked to reducing conditions such as mangrove swamps, as well as hypoxic stress in some teleosts, as ionic Mn is only available in significant quantities in the water column due to a redox reactions in low oxygen environments. Changes in Mn/Ca has also been shown to be related to fish condition in some studies.

Strontium isotope ratio (Figure 3A) exhibited a convex pattern in the sub-adult tooth, starting below 0.708 in the earliest growth periods, before increasing in the middle of the fish’s life toward the global marine signature (0.70918), then gradually decreasing to near 0.707 at the base of the tooth. In the embryo, ^87^Sr/^86^Sr was very steady and near the global marine signature of .70918 (Figure 3A).

The Sr/Ca was relatively steady in the sub-adult fish, with slightly higher values in the earliest period of growth and at mid-life (Figure 3B). In comparison the Sr/Ca was significantly elevated in the embryo, exhibiting lower values at the start of growth, increasing to their highest values between 5 and 10 mm from the tip, and decreasing slightly at the tooth base (Figure 3B).

In the sub-adult tooth, Ba/Ca exhibited the highest values at the earliest growth stages, decreasing at mid-life, and increasing again toward the end of life at the tooth base (Figure 3C. The embryo tooth contained very low levels of barium throughout, contrasting with Sr/Ca which showed the opposite pattern as would be expected in the case of movement across salinity (Figure 3C).

Magnesium to calcium ratios showed pronounced, repeated peaks throughout the life of both fish, with large decreases near the tooth base (Figure 3D). Manganese to calcium ratios showed similar repeated peaks throughout the life of the fish (Figure 3E). The highest Mn/Ca values were recorded at the tip and base of each tooth. Extreme spikes in Mn/Ca near the base of the tooth were likely caused by voids in the base of the tooth, and the extreme spike at ∼12mm is likely related to contamination rather than a biological process. Repeated peaks, however, are assumed to be related to fish movement or metabolism.

## 4. Discussion

The highly endangered status of sawfishes presents challenges to researchers and conservationists (Simpfendorfer, 2000; Kyne, Carlson & Smith, 2013; Dulvy et al., 2016). Modern, long-term, studies of living sawfish in the wild are rare and are often skewed toward smaller, juvenile fish because they are most likely to be captured (Thorson, 1982; Peverell, 2008; Poulakis et al., 2011). Further, conservation status and abundance preclude most lethal sampling of sawfish, largely limiting the promise of vertebra as an aging and microchemical tool for understanding sawfish life history and provenance, as sawfish vertebra are rarely preserved (Whitty et al., 2014). Developing methods which can address questions of life-history and location with larger sample sizes, especially of adult fish, are required to understand sawfish ecology and inform conservation efforts of sawfish. Because sawfish rostra, as opposed to vertebra, have value and are often retained, there is a larger store of historical samples available for analysis (Whitty et al., 2014; Fearing et al., 2018).

If sawfish rostral teeth do indeed contain chemical records of movement and location, as shown in this study, then archived samples may allow larger sample sizes and increased investigation of life-history questions. In particular, very little is known about the ecology and life-history of large adult sawfish due to their rarity. This has been identified as a problem in understanding basic questions of sawfish growth and maturation, and also affects our understanding of their adult movement and migration patterns (Kyne et al., 2021). Current knowledge of the locations of sawfish populations is limited in many areas, with particularly little detailed understanding of nursery areas outside of a few heavily studied nursery hotspots in Australia (largetooth sawfish, green sawfish) and Florida (smalltooth sawfish) (Poulakis et al., 2011; Feldheim et al., 2017; Huston et al., 2017; Scharer et al., 2017). Feeding areas, or movement corridors for adult sawfish of all species are less well studied (Graham et al., 2020).

These questions of habitat use and movement are critical to sawfish conservation. While sawfish are legally protected in many locations, more granular understanding of their distribution and ecology is required to provide the site and area protection and resource and habitat protection noted as a needed conservation action by the IUCN (Kyne, Carlson & Smith, 2013). If provenance and life history diversity can be investigated through the use of rostral tooth chemistry, as in teleosts and other chondricthyans, opportunities for identifying provenance and habitat use with more specificity could provide more spatially explicit ecological information to guide conservation. While the catch location of a preserved rostrum provides a single data point on species presence, analysis of rostral tooth structure and chemistry could provide temporally explicit chemical information useful for identifying nearby nursery, rearing, and feeding areas, as well as lifetime movement patterns. One of the most clear-cut examples of movement data available from calcified hard parts is movement across salinity, due to the unique dynamics of strontium and barium in relation to salinity (Zimmerman, 2005; Scharer et al., 2012; Hamer et al., 2015).

Movements across salinity can be determined using a combination of ^87^Sr/^86^Sr, Sr/Ca, and Ba/Ca, whose dynamics in teleosts and elasmobranchs appear to be similar (Tillett et al., 2011; Smith, Miller & Heppell, 2013). Strontium isotope composition is well mixed in ocean water, such that all ocean water exhibits a signature of 0.70918 (Faure & Mensing, 2004). Deviations from this ratio indicate movement into fresh or brackish water. Further, barium concentrations quickly decrease as salinity increases in estuaries, resulting in commensurate decreases in Ba/Ca in hard structures, signaling the transition from fresh to brackish water (Kalish, 1990; Walther & Limburg, 2012). Strontium concentrations, and Sr/Ca in hard parts, exhibit the opposite dynamic, increasing quickly as brackish water transitions to the higher salinity of the ocean (Zimmerman, 2005).

The sub-adult sample in this study shows likely movement across fresh and brackish water throughout its life. High Ba/Ca, low Sr/Ca, and ^87^Sr/^86^Sr far below 0.70918 indicate the earliest recorded period of the sub-adult fish was spent in low salinity, likely fresh water (Figure 3A, B&C). During the middle portion of its life (15-35mm) Ba/Ca decreases and ^87^Sr/^86^Sr increases toward the global marine signature which indicates movement toward more saline water. The lack of marked increase in Sr/Ca and the fact that ^87^Sr/^86^Sr stays slightly below the global marine signature indicates the fish was in saline water, but not in fully saline water. By the end of its life the fish appears to have moved back into fresh or mildly saline water. The smaller spikes and troughs in these signals may indicate smaller-scale forays into different salinities, either through movement or through seasonal changes in salinity as reported by Sharer et al. (2012).

This result is consistent with the known life-history of largetooth sawfish, which inhabit shallow, brackish, nearshore water as well as fresh water. Further, largetooth sawfish juveniles are known to utilize freshwater habitats as nursery areas (Wueringer, Squire & Collin, 2009). These chemical results indicate an expected pattern of freshwater utilization in early life, followed by use of multiple salinities later in life.

In contrast, the embryo chemistry indicates provenance in a much higher salinity environment. Throughout its length the embryo rostral tooth exhibits high but somewhat variable Sr/Ca, extremely low Ba/Ca, and a stable ^87^Sr/^86^Sr signature very near the global marine signature. Together, these indicate that the embryo spent the entirety of the gestation in a saline environment. The variation in Sr/Ca indicates some change of salinity, but the lack of variation in Ba/Ca and ^87^Sr/^86^Sr indicates that this change was likely not into significantly lower salinity.

Of course, the environment recorded by the embryo tooth is mediated by the internal chemistry of the mother, which presents some potential complexity in interpretation. However, it has been shown in multiple species that Sr/Ca, Ba/Ca are deposited in relation to the surrounding concentration (Bath et al., 2000; Campana & Thorrold, 2001; Smith, Miller & Heppell, 2013). Similarly, ^87^Sr/^86^Sr has been shown to deposit in hard parts without biological fractionation in the body across species (Kennedy et al., 1997, 2000; Flockhart et al., 2015). Thus, it is likely that the mother inhabited saline water during some amount of the early natal period of the embryo.

To pass these signatures on to the embryo the mother’s body must have been equilibrated to a saline environment at least until formation of the egg. As seen in salmon, provisioning by the yolk sac maintains the progeny chemistry at that of the maternal signature with little change, due to relatively low ion exchange with the outside environment (Kalish, 1990; Hegg, Kennedy & Chittaro, 2018). The same may be true of the developing sawfish whose provisioning is largely from the yolk. The rate at which the mother’s chemistry would be reflected in the embryo as it equilibrates to entry into fresh water would likely depend on the degree of ion exchange with the embryo, the speed of change within the blood, and the turnover time of the body in its entirety. None of these parameters are known in detail, though plots in Smith et al. (2013) show equilibration in Ba/Ca occurring at rates consistent with a weekly equilibration period in *Urobatis halleri*.

The Mg/Ca ratio has been called the ‘chemical clock’ in some fish species due to its association with seasonal growth rates, and one controlled elasmobranch study has found limited evidence that Mg incorporation increases with condition (Limburg et al., 2018; Pistevos et al., 2019). Without additional information about sawfish in particular it is impossible to determine, but the repeated sharp peaks in this tracer are suggestive of periods of higher and lower growth (figure 3D), a relationship which merits further study. The marked decrease in Mg/Ca near the base of the tooth (30-45mm) coincides with the onset of cloudiness in the tooth matrix. This pattern also occurs in the embryo tooth, with a markedly similar shape but scaled to the size of the smaller embryo tooth. This indicates that the change in Mg chemistry may be related to some unknown process of tooth formation occurring near the root, rather than a tracer of changes in the environment.

In teleosts, otolith Mn/Ca has been shown to loosely follow concentrations in the water, however endogenous processes related to growth further mediate it’s deposition (Mohan et al., 2012; Turner, Limburg & Palkovacs, 2015). Manganese in the water column is most closely associated with reduction of Mn oxides by microbes on the substrate under anerobic conditions and is otherwise rarely found in solution, however anthropogenic inputs from sewers have been documented as well as proximity to mangroves, which is a known habitat of largetooth sawfish (Thorrold & Shuttleworth, 2000; Mohan et al., 2012; Paillon et al., 2014; Smith et al., 2016). However the kinetics of dissolved Mn are such that it is unlikely to remain dissolved for long periods before being oxidized, and therefore increased levels in rostral teeth likely indicate close proximity to reducing environments (Laslett, 1995).

Sawfish are known to inhabit shallow, nearshore habitats and mangrove swamps where anerobic decomposition occurs, and increases in Mn/Ca could indicate forays into these environments, while faster growth during these periods would result in further apparent increases. These increases do not necessarily indicate the fish are making forays through hypoxic water, but that they may be close enough to reducing environments to incorporate dissolved Mn. It is interesting to note, however, that the peaks in Mn/Ca exhibited in both the sub-adult and embryo rostral teeth (Figure 3D) rise above the threshold set by Limburg and Casini (2018) for hypoxia exposure in teleosts, and multiple studies have shown the effects of hypoxic conditions recorded in otoliths (Altenritter, Cohuo & Walther, 2018; Limburg & Casini, 2018). However, it is unlikely that the embryo, whose gestation likely occurred in ocean water according to its Sr/Ca and Ba/Ca signatures, would have been exposed to hypoxia, indicating that incorporation of Mn into rostral teeth may differ from that of otoliths.

Analysis of banding structures in the sub-adult tooth showed that the distance between the putative growth bands formed a linear relationship rather than characteristic decrease in distance with age expected from a von Bertalanffy growth function. There are two possible reasons for this. The first is that these structures form at regular intervals along the tooth and are not influenced by growth. The second is related to tooth wear. Sawfish rostral teeth are often heavily worn, as the sub-adult sample in our study was. The amount of tooth that has worn away is difficult to know, but it is possible that the only remaining growth record in our sample is the relatively linear, sub-adult portion of the growth curve. The furthest distal rostral teeth in our rostrum were also some of the most worn, indicating that future studies should target rostral teeth with less wear and curvature, elsewhere on the rostrum.

While the length-at-age relationships of sawfish are poorly understood for larger adults, comparisons of our data to existing growth curves suggest that the putative growth bands are sub-annual. Based on the SRL/TL relationship reported by Whitty et al. (2014), the sub-adult sample was taken from a 238 cm fish. Simpfendorfer (2000) predicts the sub-adult would have been ∼5 years old, or ∼12 years old based on an alternative growth curve assuming a halved growth rate. More recent growth curves from Peverell (2008) all predict ages less than 6 years old. Thus, if the structures observed are related to age or growth, the relationship would need to be established through a relationship to other age information such as counts of annual vertebral bands.

Taken together this study presents compelling evidence that largetooth sawfish rostral teeth are a recorder of chemical information throughout the life of individual fish. Further, this chemical data records long-term, high-resolution, information about sawfish life-history and movement that is otherwise difficult to obtain from a live specimen. Our data further suggest that the structures in sectioned rostral teeth are periodic in nature and may be useful in determining growth, though they do not appear to be annual in nature.

This record of chemistry and movement may provide a useful tool in the conservation of largetooth sawfish, and likely other sawfish species. Hard-part chemistry in other fish species demonstrates the power of these chemical techniques to uncover important information about provenance, movement, and life-history which has been instrumental in conservation planning (Walther, 2019). When combined with isoscape techniques and other abiotic spatial data, hard-part chemistry is capable of uncovering important habitat and behavior with great specificity (Hobson, Barnett-Johnson & Cerling, 2010; Hamann et al., 2014; Smith et al., 2016; Brennan & Schindler, 2017). A reservoir of historical samples on which to apply these techniques exist because rostra are often saved (Whitty et al., 2014). These archived samples potentially contain information such as population-specific abiotic affinities, movement patterns, nursery hotspots, and adult movement and life-history. Combining chemical information with ongoing genetic efforts using archived rostra could also be synergistic, providing even greater detail in population connectivity and movement (Fearing et al., 2018). While these samples are necessarily historical, the information they contain is likely to be applicable to current populations, and in many cases where populations have declined precipitously may be the only information available. Further, currently a saved rostrum is used as often only a single data point recording presence of a species, often with poor spatial resolution. Rostral tooth chemistry could expand the data available from a single archived rostrum to include temporally explicit movement data, as well as population specific habitat and life history information that could support more specific conservation actions.

However, this study points out significant questions which must be answered to validate the use of sawfish rostral teeth as records of movement, life history, or growth. First, more must be known about the biology of chemical incorporation into rostral teeth, and the rates at which the chemistry of a new environment is reflected in rostral teeth. This is an ongoing question in the use of elasmobranch vertebral chemistry, and understandings from vertebral studies may provide answers applicable to sawfish rostral teeth as well.

Second, the rate of rostral tooth wear must be understood in order to understand how far back into the life of a fish the information in a rostral tooth extends. As teeth are worn, information from the early life of the fish is lost at the tip, while the most current data is available nearest the base of the tooth. Even without the entire life of the fish intact it is clear that important movement information is retained. Thus, it may still be possible to identify important nursery, rearing, or foraging locations based on chemical patterns in the existing tooth. However, understanding the rate of wear in relation to age could be used to put bounds on the time period and ages represented in sampled rostral teeth. Comparison of teeth in captive and wild sawfish, whose rate of tooth wear differs, could be used to examine this phenomenon. Sawfish rostral teeth that are entirely lost do not appear to regrow (Miller, 1974), but teeth have been reported to fracture to the pulp and return to a normal shape in at least one non-peer reviewed report (Clippinger, 1993). Methods for identifying these fractured and regrown teeth, in the same manner that regrown scales are excluded from teleost growth studies, may be important. The degree to which habitat specific factors increase or decrease wear may also be an important consideration.

The results reported here are necessarily preliminary given the limitations of our sample size. While our results are limited to largetooth sawfish, the similarities in morphology and formation of rostral teeth in other species indicate that our results may be useful for other species of sawfish as well. Larger samples sizes, including fish of known location and origin which can be compared to known water chemistries, are necessary to confirm our findings. Despite these limitations, our data supports further investigation of archived sawfish rostral teeth as a method of reconstructing movement, life-history, age and growth without the need to disturb extant fish. The large numbers of rostra housed at museums, academic institutions, and in private collections worldwide could potentially be employed to study elements of sawfish ecology which are currently difficult to study in extant populations.

## Acknowledgements

This study would not have been possible without the help of C. Caudill who provided access to the samples. Thanks to J. Vervoort and the RIGL lab at Washington State Univeristy for access to analytical equipment. Thanks to P. Charvet and V. Faria for independent species identification. In memory of P. Anders for his editing and unwavering support for this work. The authors have no conflicts of interest to report.

